# QTL combinations associated with field partial resistance to aphanomyces root rot in pea Near-Isogenic Lines

**DOI:** 10.1101/2025.02.28.640773

**Authors:** C. Lavaud, J-P. Riviere, P. Vetel, G. Aubert, P. Marget, G. Roullet, J.F. Herbommez, P. Declerck, M-L Pilet-Nayel

## Abstract

Aphanomyces root rot, caused by *Aphanomyces euteiches,* is one of the most-important diseases of pea (*Pisum sativum* L.) worldwide. The development of resistant varieties is a major objective to manage the disease. Consistent quantitative trait loci (QTL) controlling partial resistance were discovered from linkage mapping and genome-wide association studies. This study aimed to validate the resistance QTL effects and identify effective QTL combinations under contaminated field conditions, by exploiting Near-Isogenic Lines (NILs) carrying resistance alleles at individual or combined consistent QTL in different genetic backgrounds. A total of 157 NILs previously created were fingerprinted using 10,494 SNP markers from the GenoPea Infinium® BeadChip, which made it possible to confirm the QTL introgression sizes in the NILs. All NILs were phenotyped for resistance in field contaminated nurseries over two years at six locations in France. NILs carrying resistance alleles from PI180693 or 90-2131 at the major-effect QTL *Ae-Ps7.6*, individually or in combination with minor-effect QTL (*Ae-Ps4.1* or *Ae-Ps5.1*), showed significantly increased levels of partial resistance in different environments and genetic backgrounds. At other QTL combinations (*Ae-Ps1.2 or AePs7.6 + Ae-Ps2.2 + Ae-Ps3.1*), alleles from PI180693 or 552 also showed significant effects on partial resistance in some NIL genetic backgrounds. At these QTL combinations, the PI180693 resistance alleles also contributed to late flowering. This study provides tools and information for the choice of resistance QTL to combine in breeding, to increase partial resistance to *A. euteiches* in pea varieties.

## INTRODUCTION

Polygenic plant resistances controlled by numerous genomic regions, or Quantitative Trait Loci (QTL), typically confer partial and potentially durable resistance to diseases in crop species (Cowger and Brown 2019). However, exploiting QTL knowledge in breeding programs remains challenging due to the partial levels, poorly known functions and instability of their effects, partly attributed to their interactions with the genetic background and environment (Nelson et al. 2018). Near-Isogenic-Lines (NILs) have been useful to characterize and validate the stability of QTL effects. QTL-NILs are typically produced by transferring (‘introgressing’) one or more chromosomal segments, harboring QTL confidence intervals, from a resistant genotype into the genetic background of a susceptible line. NILs can therefore be used to characterize contrasting chromosomal segments on a uniform genetic background. Most NILs are created through a few generations of backcrossing with a single ‘recipient’ parental line to introgress fragments from a ‘donor’ line. The development of genomic technologies has made it possible to better characterize plant genomes and develop tools such as Single Nucleotide Polymorphism (SNP) marker arrays (Thudi et al. 2021), to better characterize chromosomal segments and especially disease resistance QTL in NIL-type material (Qiu et al. 2022).

Aphanomyces root rot of pea (*Pisum sativum L.*), caused by the soilborne oomycete *Aphanomyces euteiches* (Jones and Drechsler 1925) has become one of the most damaging diseases of peas worldwide since the 1990’s. It can cause yield losses up to total crop loss in a highly contaminated field (Pfender et al. 2001). *A. euteiches* has been especially reported in the US (Gossen et al. 2016), Canada (Wu et al. 2018), France (Quillévéré-Hamard et al. 2018), Australia (van Leur et al. 2008) Sweden (Levenfors et al. 2003) and the Netherlands (Oyarzun and van Loon 1989). Symptoms appear at a very early plant stage (from 3 to 4 leaves) with soft, translucent lesions on the rootlets, which progress into a brown-honey root affecting the entire root system up to the epicotyl. Root damages result in yellowing, stunting, and possible drying of plants on the aerial parts. Two main pathotypes of *A. euteiches* have been reported in pea based on their differential reactions on a set of six genotypes (Wicker and Rouxel 2001; Wicker et al. 2003). Pathotype I is predominant in Europe and has been observed in the USA and Canada. Pathotype III has so far only been reported in certain regions of the USA and Canada (Sivachandra Kumar et al. 2021; Moussart et al. 2024). No efficient chemicals are available to control the disease. Genetic control, combined with the use of prophylactic and cultural methods, is currently the most promising solution to manage the disease. In pea, resistance to *A. euteiches* exhibits a partial level and is polygenically inherited (Pilet-Nayel et al. 2002). Since the early-2000s, genetic dissection of polygenic resistance to *A. euteiches* has been perdormed by linkage mapping in recombinant inbred line (RIL) and Advanced Backcross (AB) populations, as well as by Genome-Wide Association Study in a Pea-Aphanomyces collection (Pilet-Nayel et al. 2002; Hamon et al. 2011; 2013; Desgroux et al. 2016; Wu et al. 2021; Leprévost et al. 2023). Comparative genetic analysis of resistance across these populations using SNPs identified ten consistent genetic regions and resistance haplotypes cumulated in some germplasm that were useful as sources of resistance for breeding. Especially, seven genetic regions were consistently detected from multiple populations, environments, and/or different strains of *A. euteiches*. These regions included two major-effect QTL, *Ae-Ps7.6* and *AePs4.5*, as well as the minor-effect QTL *Ae-Ps1.2*, *Ae-Ps2.2*, *Ae-Ps3.1*, *Ae-Ps4.1* and *Ae-Ps5.1*. At some of them, linkages were identified between resistance and undesired alleles for feed pea breeding (*i.e.* late flowering at *AePs2.2* and *Ae-Ps3.1*; anthocyanin production at *Ae-Ps2.2;* normal leaves at *Ae-Ps1.2*), especially when brough by the partially resistant germplasm PI180693. A Marker-Assisting Backcrossing (MAB) program was implemented to simultaneously introgress resistance alleles at one, two, or three of these seven consistent genetic regions, from five donor lines into three or four recipient lines, using Simple Sequence Repeat (SSR) markers (Lavaud et al. 2015). This program resulted in the development of 157 pea NILs, carrying resistance alleles at none, one, two, or three resistance QTL in different genetic backgrounds. The evaluation of these NILs under controlled conditions validated the two major-effect QTL (*Ae-Ps7.6* and *Ae-Ps4.5*) against *A. euteiches* strains from pathotypes I and III, respectively (Lavaud et al. 2015; Lavaud et al. 2024). It also pointed out QTL x genetic background interactions, particularly at the major QTL *Ae-Ps7.6*. However, further knowledge was required regarding the QTL effects in NILs in field contaminated environments as well as on the fine characterization of the genomes of the created NILs. This knowledge would be relevant to validate the effect of QTL combinations on increasing the level of partial resistance and support strategies for choosing best QTL combinations for breeding.

The objective of this study was to validate the effect of QTL in NILs, individually or in combination, on partial resistance to Aphanomyces root rot in field conditions, regarding their fine genomic composition. The 157 NILs were phenotyped for resistance on root and aerial plant parts in six field contaminated nurseries over two years. The NILs were genotyped using the GenoPea Illumina Infinium® BeadChip (Tayeh et al. 2015), to finely characterize the genomic regions inside and outside the chromosomal segments introgressed and thus identify the contribution of the genotypes to the observed phenotypic variations. The results validated the effect of several QTL combinations, consistently or specifically, on partial resistance in different environments and plant organs affected by the disease in the field. They identified most effective combinations of QTL to limit disease severity.

## MATERIAL AND METHODS

### Plant material

The 157 pea NILs carrying resistance alleles at none, one, two, or three resistance QTL to *A. euteiches*, described in Lavaud et al. (2015), were used in this study. The NILs were derived from biparental crosses between donor lines (RIL 831.08, RIL 847.50, RIL BAP 8.70, RIL BAP8.195, 552) and recipient lines (reference line, *i.e.*, Baccara or DSP or Puget; spring pea variety Eden, winter pea variety Isard and Enduro) (Lavaud et al. 2015). The donor lines originated from the pea germplasm 90-2079, 90-2131 (Kraft 1992), PI180693 (Lockwood and Ballard 1960), and 552 (Gritton 1990), which were used as resistant parents of the RIL mapping populations previously analyzed for QTL detection (Hamon et al. 2013; Lavaud et al. 2015). The recipient reference lines were the susceptible parents of the RIL mapping populations. The 157 NILs were derived from 16 parallel backcrossing schemes resulting in 16 sets of NILs, each NIL set comprising two sister NILs (named a, b or c) containing the same QTL or QTL combination. Genotyping and field phenotyping of the NILs were performed using plants at BC5/6F2-3 and BC5/6F3-5 generations, respectively.

A total of 14 control lines were used in the field resistance tests, including the 11 parental lines of the tested NILs and three control lines with high levels of resistance (Roux-Duparque et al. 2004; Desgroux et al. 2016). Among the three resistant controls, AeD99OSW-45-8-7 and AeD99OSW-50-2-5 were selected from crosses between three of the four resistant parental lines of the QTL mapping populations previously studied [(90-2131xPI180693) x 552], and AeD99QU-04-9-10-1 was obtained from the cross between four lines [(Baccara x PI180693) x (Capella x MN314)] (Desgroux et al. 2016).

### Genotyping of NILs

The DNA of all NILs was extracted using a NucleoSpin Plant II extraction kit (Macherey Nagel GmbH & Co., Germany), then quantified by fluorescence (PicoGreen) on the PheraStar, and finally normalized to 100ng/μl. Genotyping of the NILs was performed using the GenoPea Infinium® BeadChip array (ANR-GenoPea project), which includes 13.2K SNPs located in gene sequences and associated with distinct transcripts (Tayeh et al. 2015). The SNPs were analyzed individually for each of the 16 NIL sets using version 1.9.4 of the Illumina GenomeStudio® software module. A consensus map of 15,079 markers, including 12,802 SNPs from the GenoPea chip, was previously constructed from 12 differents RIL populations (Tayeh et al. 2015). The ordering of markers from this consensus map and the genotyping data of the NILs were used to generate graphical genotypes of the NILs.

### Evaluation of NILs in field contaminated nurseries

The 16 NIL sets and the control lines were evaluated for resistance to *A. euteiches* in the field on a multi-local network of six highly contaminated nurseries in France, located at (i) Riec-sur-Belon (RB) – Finistère, Bretagne (UNILET-INRAE; (Hamon et al. 2011); (ii) Dijon-Epoisses (DI) – Côte d’Or, Bourgogne-Franche-Comté (INRAE; (Hamon et al. 2011), (iii) Froissy (FO) – Oise, Haut de France (Unisigma), (iv) Mons-en-Pévèle (MP) – Nord, Haut de France (KWS-Momont) and (v) Louville-la-Chenard (LC) and Chartainvilliers (CH) - Eure-et-Loir, Centre-Val de Loire (RAGT2n and Limagrain Europe, respectively). The lines were evaluated over two years, 2014 and 2015, at all sites.

The field experiments were conducted in a randomized complete block design, with three blocks in the RB and DI nurseries and two blocks in the other nurseries. Each plot in each block consisted of a double row of two meters long, with each row containing 30 seeds. The design included the spring pea variety Solara as an adjacent susceptible control every four plots, to control for heterogeneity of soil contamination within the nursery.

All the pea lines were evaluated for resistance using two traits. (i) Disease severity on roots (DS) was measured from the four-leaf stage on ten plants per plot. Each plant was uprooted keeping the roots intact, then washed in water and scored using a 0 to 5 scale, adapted from (Moussart et al. 2001), with 0 for a healthy plant, 1 for translucent to beige traces on the rootlets, 2 for beige to brownish, soft areas covering a quarter of the root system and the epicotyl being healthy, 3 for beige to brownish, soft areas covering at least half of the root system and the epicotyl may turn beige but remains firm, 4 for a root system attacked three-quarters and the brownish epicotyl is soft and appears constricted, and 5 for a dead plant. This scale is similar to the one previously used by Lavaud et al. (2015) for NIL evaluation under controlled conditions. (ii) The aerial decline index (ADI) was evaluated on each plot twice every 10-15 days from the late flowering plant stage. A scoring scale from 1 to 8, as in Hamon et al. (2011) and adapted from Duparque and Boitel (2001), was used in all nurseries. This scoring considers plant yellowing.

All the pea lines were also evaluated for two developmental traits, including earliness at flowering (FLO) and plant height (HT). FLO was scored on each plot as the number of days to 50% bloom from the first day of the year. HT measured the average height of five plants at maturity in a whole plot. For all NILs, HT and FLO were measured within one block in healthy nurseries at four of the network sites (FO, MP, LC, CH) and at Le Rheu (LR) - Ille-et-Vilaine, Bretagne (INRAE).

### Statistical Analysis

Statistical analyses of the phenotypic disease (DS, ADI) and developmental (HT and FLO) scoring data were conducted using R software (R Core Team 2020). The analyses used individual plant scores for the DS variable and overall plant scores by plot for the other variables (ADI, HT and FLO). For each variable, global multi-environment and individual environment data analyses were performed, firstly on all evaluated lines and secondly on each NIL set, using a linear mixed model (LMM; lmer function from the ‘lme4’ package; (Bates et al. 2015)) and a deviance analysis to estimate the effects of the genotype, location, year and block (Wald test, α=5%; Anova function from the ‘car’ and ‘RVAideMemoire’ packages;(Fox and Weisberg 2019; Hervé 2020)). A principal component analysis (PCA) was performed to estimate the correlations between the developmental (HT, FLO) and resistance (DS and ADI) variables from the multi-environment or individual analysis (‘FactoMineR’ package; (Lê et al. 2008)).

The global analysis performed on all NILs considered the genotype, location, year and interactions as fixed factors and the block as a random factor, to estimate the effect on the observed score variations. The global analysis performed on each NIL set only considered the genotype as a fixed factor and the location, year and block as random factors. For each NIL or NIL set and each variable, an adjusted mean of the location, year and block effects was estimated from the analysis (LSMeans; “lsmeans” function from the “lsmeans” package; Searle et al. 1980; Lenth 2023). The LSMean values obtained for the NILs within each set were compared with each other, using a Tukey multiple comparison test (α=5%; ‘cld’ function from the ‘MultcompView’ package; (Luciano 2006)). Broad-sense heritability (h²) was calculated for each variable from the multi-environment analysis according to the formula h² = σ²g / [σ²g + (σ²e / n)], where σ²g is the genetic variance, σ²e is the residual variance, and n is the number of environments, from a Wald F test considering all NILs (Halekoh and Højsgaard 2014).

The individual analysis by environment conducted on all NILs or on each NIL set, used the genotype as a fixed factor and the block as a random factor. For each NIL and variable, an adjusted mean of the block effect was estimated from the analysis. Within each NIL set, LSMean values of NILs carrying QTL were compared with the average of sister NILs without, using the pairwise contrast test (pairs’ function). Mean contrast values over the different environments was calculated for each QTL-carrying NIL and each DS or ADI variable.

## RESULTS

### NIL graphical genotypes

Among the 12,802 SNP markers of the GenoPea chip mapped by Tayeh et al. (2015), 10,494 were polymorphic in at least one of the 16 NIL sets studied and/or the four pairs of parental lines of the RIL populations previously used for QTL detection (Hamon et al. 2013). Between 1,040 and 3,181 polymorphic SNPs between the parental lines of the donor RILs made it possible to locate regions presenting the same genome as that of the reference recipient lines, in the donor RILs of the BAM program and subsequently the NILs obtained on reference backgrounds (Figure 1). In addition, between 1,726 and 4,795 polymorphic SNPs between the parental lines of each NIL set, depending on the NIL set, allowed the characterization of the NIL genomes (Figure 1). The number of polymorphic SNPs between the donor parental lines and the Eden recipient line was 200 to 1000 SNPs lower, depending on the crosses, compared to that between the donor lines and the Isard recipient line. Out of the genotyping points used to construct the graphical genotypes of the NILs, 0.6% revealed heterozygous genotypes, most of which were located in QTL flanking regions or outside QTL regions (Figure 1).

**Figure 1:**
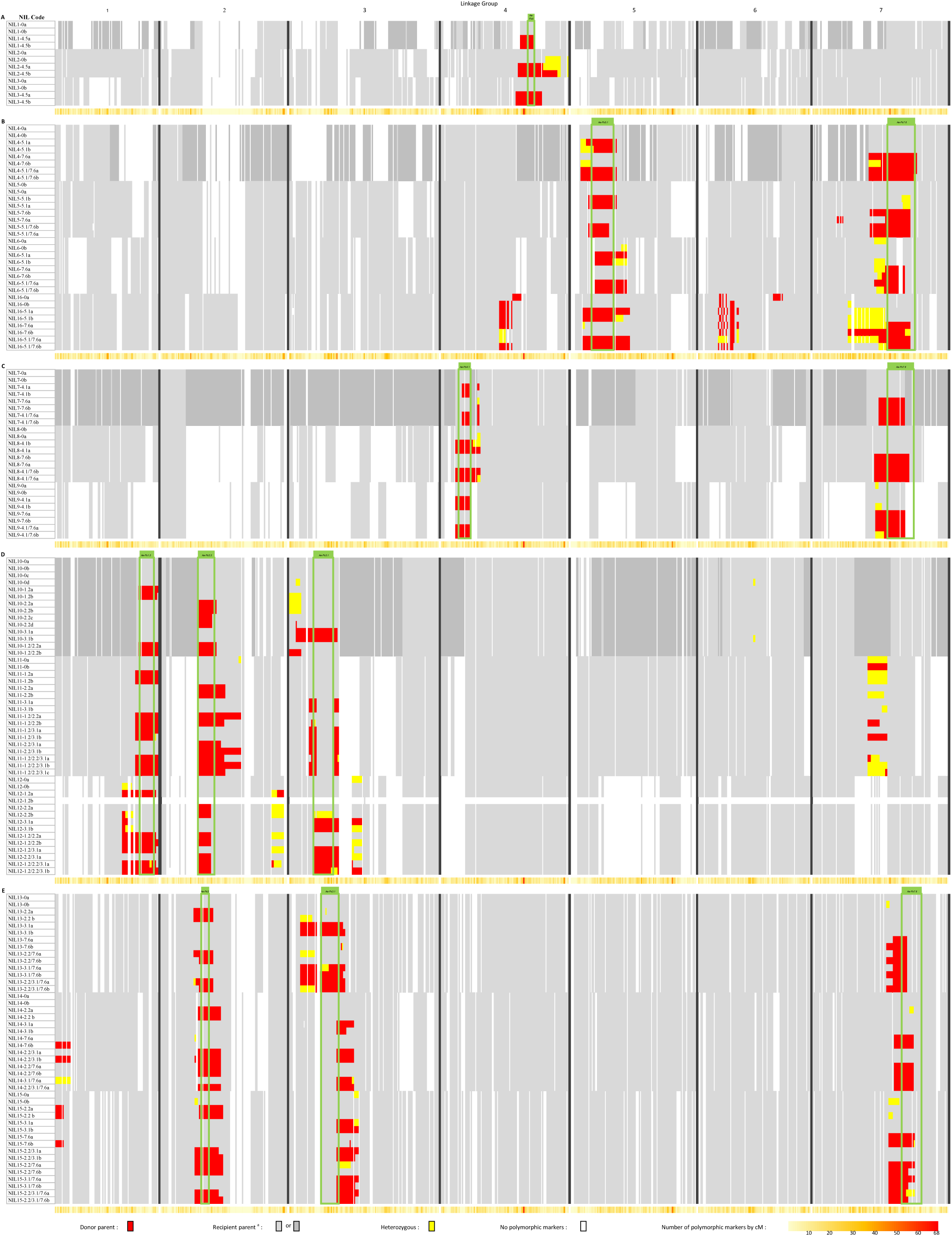
Genomic fingerprint of the 157 pea NILs at zero, one, two or three of the seven main QTL for resistance to *A. euteiches* selected in the MAB program from (Lavaud et al. 2015). The white, dark grey, light grey, red and yellow colors indicate, respectively, for each NIL: non-polymorphic markers; polymorphic markers between the parents of the donor RILs and revealing the presence of the reference recipient genome (Baccara, Puget or DSP) in donor RILs; polymorphic markers between the parents of NILs showing a return to the recipient genome; and polymorphic markers between the parents of NILs revealing a donor fragment at homozygous or heterozygous state. Green boxes show confidence intervals of resistance QTL targeted for introgressions, according to (Leprévost et al. 2023).

On the linkage groups (LG) carrying QTL, the graphical genotype analysis identified variable introgression sizes of the donor fragments, either due to variable level of return to the recipient genome in the QTL flanking regions, or to loss of donor segments in the targeted QTL regions (Figure 1). Nearly all NIL sets have introgressions larger than the confidence intervals (CI) of the target QTL. For example, for the NIL4 set, the introgressions at both QTL *Ae-Ps5.1* and *Ae-Ps7.6* are twice as large as the CI of QTL (Figure 1). Three NIL sets (NIL11, 14 and 15) have lost introgression into the center of QTL *Ae-Ps3.1* and have retained parts of the donor segments in one and/or the other flanking regions of the QTL. Some NILs [NIL12-1.2/2.2/3.1(a and b) and NIL13-3.1/7.6a] showed heterozygosity at markers in the target QTL regions. Several NILs were also heterozygous in the flanking regions of the target QTL [for example, in NIL6 set at QTL *Ae-Ps7.6*, NIL4 and NIL6 sets at QTL *Ae-Ps5.1*, NIL8 set at QTL *Ae-Ps4.1*] or in QTL regions not selected in NILs [for example, NIL5-5.1(a and b) and NIL14-2.2a at QTL *Ae-Ps7.6*].

The graphical genotypes of the NILs showed a level of return to the recipient genome outside the QTL regions exceeding 90% for all NILs (data not shown). Some donor fragments, in homozygous or heterozygous states, remain present outside the QTL regions, especially in the NIL sets resulting from schemes aiming to introgress three QTL simultaneously. Thus, residues of donor fragments were present on LGI and LGVII not carrying target QTL in NIL14 and NIL11 set, respectively. The same observation was made on LG carrying QTL in regions neighboring or not of target QTL, *i.e.*, in the sets NIL16 on LGVII, NIL10 on LGIII and NIL12 on LGI, II, and III (Figure 1).

Despite the dense genotyping of the NILs, not all genome regions have been covered by polymorphic markers (Figure 1). The absence of polymorphism between NIL parental lines in these regions, sometimes of large size (65 cM on LGII, 52 cM on LGV), was found in regions where the donor parental line had the same genome as that of the reference recipient parent, or even as other recipient varieties.

### Evaluation of NILs for resistance to Aphanomyces root rot in field conditions

The analysis conducted on the adjacent Solara control repeated once every four plots in the experimental design highlighted significant heterogeneity in soil inoculum potential within the FO, MP, LC and CH nurseries, but not within the RB and DI nurseries, over the two years (Additional file 2). Therefore, analyses of DS data obtained at RB and DI were performed using root rot scores on individual plants, unweighted by adjacent control scores. Analysis of ADI data obtained at all sites were conducted using overall plant scores per plot weighted by those of adjacent controls. ADI data were retained for analysis in seven of the 12 experiments, due to inconsistent control distributions or low disease levels in the other experiments. For the HT and FLO variables, data were obtained for analysis in five and four experiments, respectively (Additional file 3).

Global multi-environment data analysis on all NILs in the retained experiments showed highly significant genotype, location and year individual and interaction effects (P<0.001) for the four variables (DS, ADI, HT, and FLO). Individual data analysis by environment on all NILs revealed a highly significant genotypic effect (P < 0.001) in each experiment for each variable. The heritability of the traits (H²), estimated from the multi-environment analysis for DS, ADI, FLO, and HT variables, was moderate to very high, with values of 0.82, 0.79, 0.96, and 0.54, respectively. Global LSMean values for DS and ADI obtained for all NILs showed a continuous and Gaussian distribution (Figure 2A, Additional file 3). The pea control lines ranked as expected, with the parental donor lines of NILs showing moderate to high levels of partial resistance and the resistant controls (Roux-Duparque et al. 2004) showing the best levels of partial resistance. Transgressive NILs, *i.e.* more susceptible or more resistant than the parental lines, were identified (Figure 2A). However, no NILs were more resistant than the AeD99OSW-45-8-7 or RIL 847.50 controls for root or aerial resistance, respectively (Figure 2A, Additional file 3).

**Figure 2:**
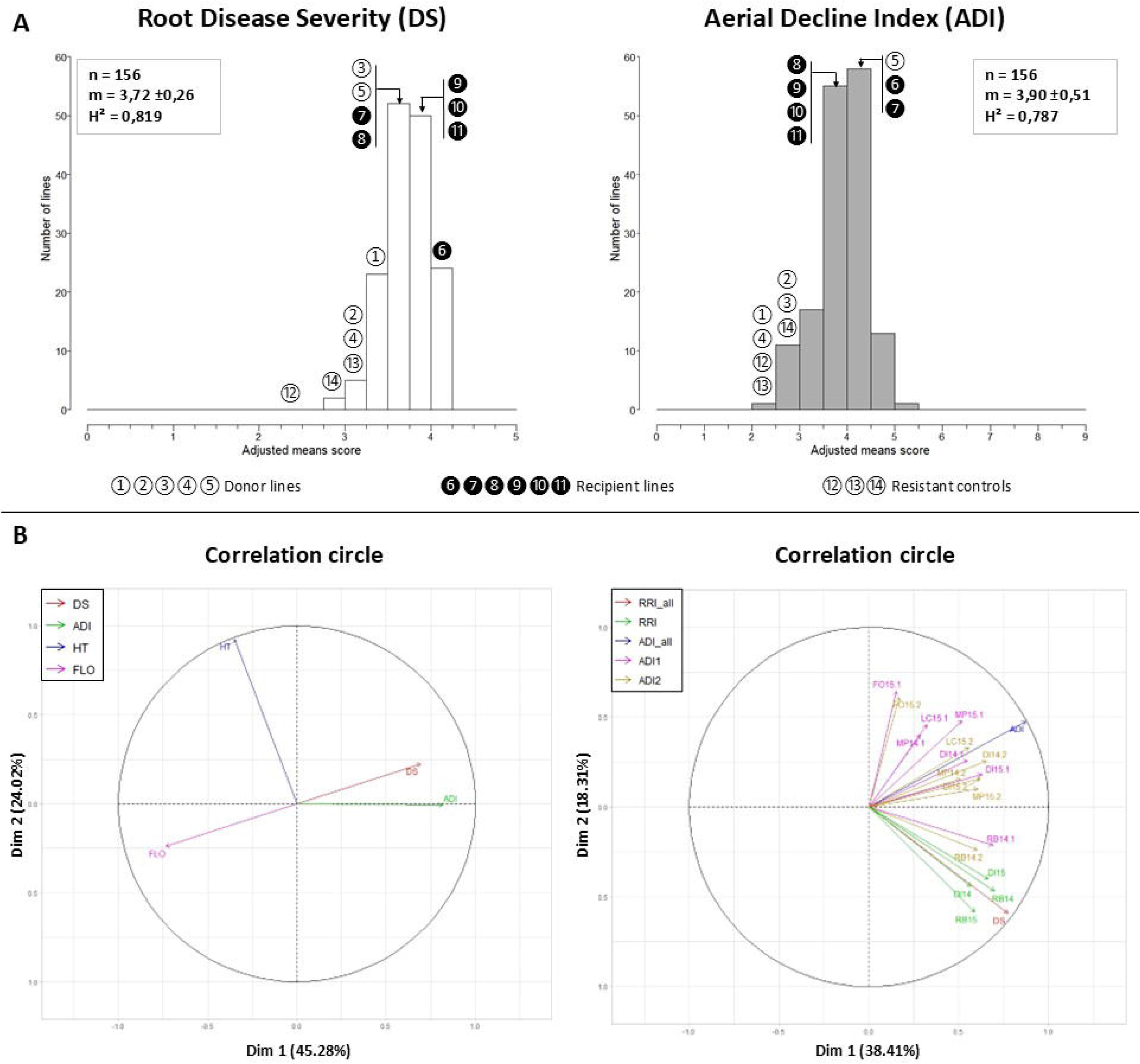
Frequency distribution of field disease values caused by *A. euteiches* and correlations between variables, obtained on 157 pea NILs over six sites and two years in contaminated nurseries. A-Frequency histograms of the root disease severity (DS) and aerial decline index (ADI) values of the 157 pea NILs. The parental lines of “donor” and “recipient” NILs are shown in white and black, respectively: 1-RIL 847.50, 2-RIL BAP8.195, 3-RIL BAP8.70, 4-552 and 5-RIL 831.08; 6-Eden, 7-Isard, 8-DSP, 9-Puget, 10-Baccara and 11-Enduro. The values of the resistant controls are shown in white: 12-AeD99OSW-45-8-7, 13-AeD99OSW-50-2-5 and 14-AeD99QU-04-9-10-1. n: total number of NILs evaluated, m: mean standard deviation of all NILs, H²: heritability. B-Circle of correlations for the different variables studied (DS: root Disease Severity; ADI: Aerial Decline Index; FLO: Flowering; HT: plant Height), and the different experimental sites (DI: Dijon-Epoisses; FO: Froissy, LC: Louville-la-Chenard; MP: Mons-en-Pévèle; RB: Riec-sur-Belon) and years (2014, 2015).

The PCA analysis from the global LSMean values indicated a negative correlation between root or aerial symptoms and flowering time (r²=-0.3 and −0.44, respectively), suggesting that resistant lines tend to flower later. No correlation was identified between plant height (HT) and the other variables (−0.21<r²<0.08), indicating that the introgressed resistance QTL did not affect plant height (Figure 2B). High correlations between sites were observed for DS values, whereas correlations were more variable between sites for ADI values, with some being zero (*e.g.*, RB14 and FO15) and others being high (DI14, DI15, and LC15) (Figure 2B).

### Effects of individual and combined QTL on field resistance within NIL sets

Consistent significant differences between NILs with and without QTL in sets containing resistance alleles from the donor lines 90-2131 (RIL 847.50) or PI180693 (RIL BAP8-70 and RIL BAP8-195) were identified for DS and ADI variables, using multiple comparison tests of multi-environment LSMean values (Additional file 4) and means of the contrast values obtained in the different environments between NILs with and without QTL (Table 1), within each NIL set.

**Table 1:**
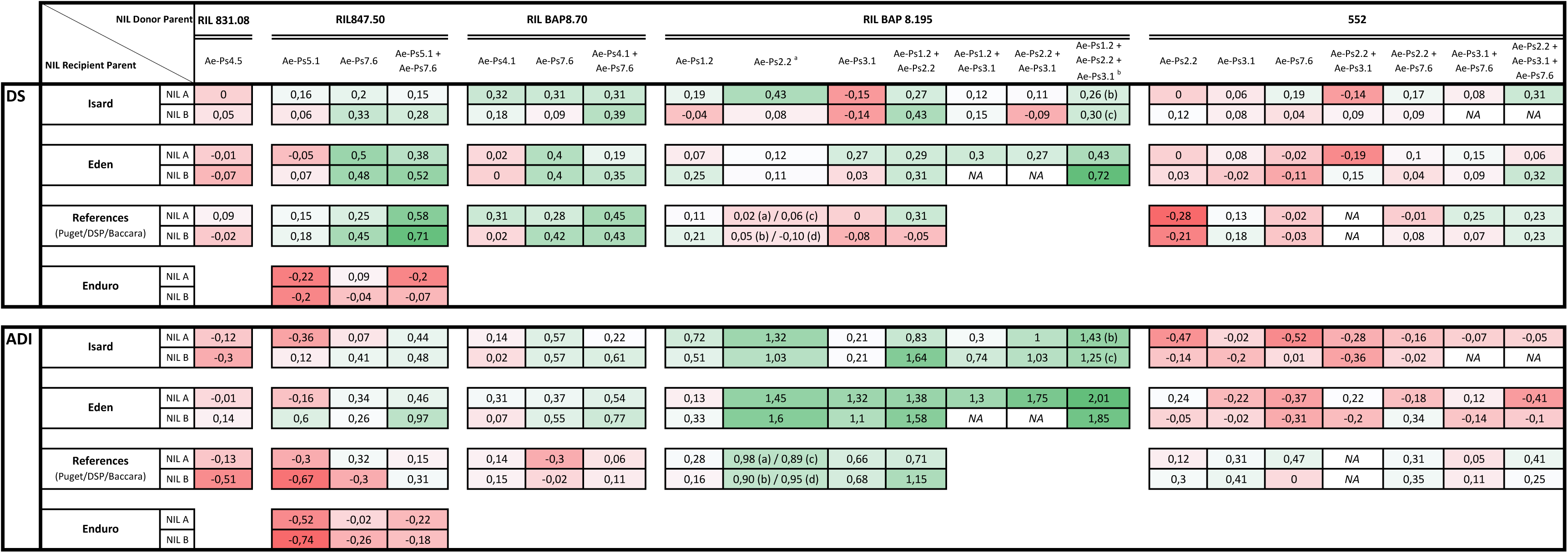
Mean contrast values between NILs with and without QTL, for disease severity on roots (DS) and aerial decline index (ADI) in all location*year trials, by donor line and QTL combination in the different genetic backgrounds. Contrast values were obtained in each NIL set as follows: the multi-environment LsMean DS or ADI values of the QTL-carrying NILs were subtracted from the average values of the two sister NILs without QTL (NILx-0a, NILx-0b). A color scale from red to green represents a resistance level of a QTL-carrying NIL from lower to higher, respectively, than that of the QTL-free NIL from the same set. ^a^ Four sister NILs carrying QTL *Ae-Ps2.2* obtained in the « Baccara » genetic background are mentioned by the letters a, b, c and d. NA: missing data.

In NILs carrying introgressions from RIL 847.50, variable effects of QTL *Ae-Ps7.6*, alone or combined with *Ae-Ps5.1*, were observed depending on the variable (DS or ADI) and genetic background. (i) For DS, significant effects of *Ae-Ps7.6*, individually or in combination with *Ae-Ps5.1*, were observed in one to three environments depending on the genetic background (Additional file 5a). Global contrast analysis of DS values revealed a significant effect of a NIL (NIL4-5.1/7.6b) carrying the combination of the two QTL in the reference background compared to NIL without QTL. (ii) For ADI, only NIL6-5.1/7.6b showed a significant effect on resistance, with a notable mean gain of 0.97 over the environments tested (Table 1). For QTL *Ae-P5.1* individually, contrast analysis showed no significant effects on DS or ADI. No significant QTL effects were observed in the Enduro background for either DS and ADI variable. For morphological variables (HT and FLO), no significant QTL effects were detected in any genetic background.

For NILs carrying introgressions from RIL BAP8.70, contrast analysis with NILs lacking QTL showed a significant individual effect of QTL *Ae-Ps7.6* on DS in different environments in the three genetic backgrounds, especially Baccara (Additional file 5a, Table 1). This effect was also observed for ADI, with a higher value in the winter Isard background compared to the reference background. NILs carrying QTL *Ae-Ps4.1* individually exhibited a significant effect on DS in two genetic backgrounds, Baccara and Isard, each NIL in a single environment. Some of these NILs also exhibited an effect on ADI in some environment, especially in Eden (Additional file 5b). The combination of the two QTL showed significant effects in one or two environments, for DS in all NILs and for ADI in NILs in Eden with a mean gain of 0.77 (Additional file 5a, Table1). No significant effects were observed between NILs for the two morphological variables, except for NILs carrying the two QTL in Baccara and Eden for FLO.

In NILs carrying introgressions from RIL BAP8.195, significant effects of QTL *Ae-Ps3.1* and *Ae-Ps2.2*, individually or in combination, were observed in Eden background, on ADI and DS in one or two environments, as well as on flowering time. In the two other genetic backgrounds, Baccara and Isard, QTL *Ae-Ps2.2*, individually or in combination with *Ae-Ps1.2*, had significant effects on ADI and flowering time (Additional file 4). Genotyping analysis of NIL11 set also revealed introgression of a donor region upstream of the QTL *Ae-Ps7.6*. This introgression of the Baccara allele into NILs in the Isard background (NIL11-0b, NIL11-1.2/2.2b, and NIL11-1.2/3.1b) was associated with a negative effect on flowering time (Additional file 4). Contrast analysis also identified significant effects of QTL *Ae-Ps1.2* and *Ae-Ps2.2*, individually and/or in combination, for ADI in one or two environments and at least one recipient background (Additional files 5a and 5b). Only one NIL carrying the combination of three QTL (*Ae-Ps1.2*, *Ae-Ps2.2*, and *Ae-Ps3.1*) was identified in the Eden background, since introgression at *Ae-Ps3.1* was not validated in Isard background based on genotyping data. This NIL (NIL12-1.2/2.2/3.1b) demonstrated consistent significant effects on reducing both DS and ADI scores, with mean gains of 0.72 and 1.85, respectively, compared to the QTL-free NILs (Table 1).

Among the NILs carrying introgressions from the donor line 552, no significant differences in mean ratings were observed for any variables (Additional file 4). The QTL *Ae-Ps2.2* and *Ae-Ps3.1* did not show effects on flowering time, unlike their effects observed in NILs derived from the donor line RIL BAP8.195. Contrast analysis by environment showed significant environment-specific effects of the three QTL *Ae-Ps2.2*, *Ae-Ps3.1*, and *Ae-Ps7.6* in combination across all three genetic backgrounds, but only for DS.

In NILs carrying introgressions from RIL 831.08, no significant effect of the QTL *Ae-Ps4.5* was observed on the different variables, except in a single environment for DS and/or ADI in the spring genetic backgrounds Puget and Eden (Additional file 5a).

## DISCUSSION

This study first reports the fine whole-genome characterization and field evaluation for Aphanomyces root rot resistance of a large number of pea NILs with introgressions carrying resistance alleles from different sources of resistance to *A. euteiches*. Few studies have finely characterized the whole genome of NILs produced to validate QTL effect or analyze their underlying mechanisms(Ando et al. 2008; Schmalenbach et al. 2008; Ding et al. 2011; Navara and Smith 2014; Badis et al. 2015).

### Genomic characterization of NILs

Genotyping of the 157 NILs using 10,494 SNPs, of which 27% to 48% were exploited to characterize each NIL, confirmed the presence of introgressed segments, with variable sizes depending on the NILs, except at the QTL *Ae-Ps3.1* (allele from 552 or PI180693) in three sets of NILs. It also identified genomic regions outside the QTL that present residues of donor fragments, especially in the QTL flanking regions, or that remain unassigned to parental genomes due to lack of polymorphic markers. Compared to the previous description of the NIL genome (Lavaud et al. 2015), this dense genotyping of all NIL sets provided more precision to confirm their genomic content.

In most of the NILs, the size of introgressed fragments was estimated to be larger than the confidence interval (CI) of the previously detected QTL (Pilet-Nayel et al. 2002; 2005; Hamon et al. 2011; 2013; Leprévost et al. 2023). On one hand, the low number of available markers (SSRs) at the time of NIL construction led to less stringent and less precise selection of individuals at the QTL as well as in the flanking regions. On the other hand, the number of plants used in each MAB cross and generation during NIL production, *i.e.* 45 to 91 plants depending on the number of simultaneous introgressions (Lavaud et al. 2015), did not allow for strong selection pressure on the QTL flanking regions, as discussed in Lavaud et al. (2015). Nevertheless, the dense genotyping of NILs validated the presence of donor alleles at the QTL CI within the introgressed fragments, in 12 of the 16 NIL sets.

Losses of donor fragments in QTL regions were observed in four NIL sets. Introgressions from donor parents were lost at the QTL *Ae-Ps3.1* as suggested by Lavaud et al. (2015), as well as at the QTL *Ae-Ps7.6* for part of the introgression. These lost fragments were mainly due to the lack of polymorphic markers available to trace introgressions along their entire lengths. In NIL set 11, none of the three SSR markers used to trace the introgression from PI180693 at QTL *Ae-Ps3.1* were located in the central part of the QTL CI. This resulted in the loss of the donor fragment, likely by double recombination in NIL set production process. In total, 23 NILs lacking the introgression at QTL *Ae-Ps3.1* were therefore not eligible for QTL validation, individually or in combination. At QTL *Ae-Ps7.6*, part of the introgression was lost in NIL sets 13 and 14, but this did not result in differences in phenotypic effect compared to NILs carrying the introgression in NIL set 15. Hospital et al. (2005) advocated the use of at least three markers per CI of a QTL, including two flanking markers and one marker in the middle of the CI, to be able to successfully introgress a QTL in MAB schemes.

The presence of donor fragments outside the introgressed region may bias the evaluation of the QTL effect. In our study, we observed a very good return to the recipient parent genomes after five or six generations of backcrossing to the recipient backgrounds. Nevertheless, some residues of donor genome fragments were identified on LGI, II, III outside the QTL regions and their flanking regions, especially in NIL sets derived from introgression schemes of three QTL simultaneously. In these schemes, the number of plants tested at each generation (n=91) did not allow for a strong selection rate outside QTL regions. Regions with donor genome residues were located outside QTL regions not targeted for introgressions, except at QTL *Ae-Ps7.6* in certain NILs from the set 11 (*e.g.* NIL11-2.2b) but these residues did not generate biais on the phenotype.

### Individual QTL effects in pea NILs on field partial resistance

NIL phenotyping data obtained for field resistance on root and aerial plant parts validated, in multiple recipient backgrounds and/or environments, the individual effects of (i) one of the two major-effect QTL, *Ae-Ps7.6*, (ii) the QTL *Ae-Ps4.1* that was previously unvalidated in NILs under controlled conditions and (iii) the QTL *Ae-Ps2.2* and *Ae-Ps5.1*, previously observed in NILs under controlled conditions.

The effect of the major QTL *Ae-Ps7.6* in pea NILs, which was observed under controlled conditions with a high level on root resistance to *A. euteiches*, was confirmed for field resistance on the roots, with variable levels depending on the introgressed allele. The effect of the 90-2131 and PI180693 alleles were significantly highlighted in at least one pea NIL genetic background. The effect of the 552 allele at the QTL *Ae-Ps7.6* in pea NILs was not significant in reducing root disease severity in the field, although it was observed previously under controlled conditions. These field observations are consistent with previous QTL detection data (Hamon et al. 2011; Leprévost et al. 2023), which reported that the effect of *Ae-Ps7.6* was consistent across different soils and experimental years. In our study, the effect of *Ae-Ps7.6* on reducing root disease severity in the field was more consistent in spring than in winter recipient backgrounds, contrary to observations made under controlled conditions by Lavaud et al. (2015). However, *Ae-Ps7.6* showed no significant individual effect on reducing aerial disease symptoms in the highly contaminated nurseries and NIL genetic backgrounds studied, while it was consistently detected for ADI variables in previous QTL studies (Hamon et al. 2013; Leprévost et al. 2023).

The QTL *Ae-Ps4.5*, whose major effect was detected under controlled conditions by Lavaud et al. (2015) with the Ae109 strain (reference for pathotype III - USA), showed a significant effect in Dijon in 2014 and/or 2015 in at least one NIL genetic background. To date, no *A. euteiches* isolate from pathotype III has been identified in France. However, Moussart et al (2024) identified three isolates of undetermined pathotype in the contaminated Dijon nursery used to evaluate NILs in this study, in addition to seven other isolates characterized as pathotype I. Of the three isolates, “Di9” showed a pathogenicity profile on the differential pea range close to pathotype III. In the contaminated nursery of Riec-sur-Belon (FR) used for NIL phenotyping in our study, and the one of Pullman (WA, USA), the authors characterized isolates as all belonging to pathotype I. Leprévost et al (2023) detected *Ae-Ps4.5* from aerial disease scorings on the RIL Puget x 90-2079 population in the Pullman nursery, suggesting that the QTL may not be specific to pathotype III.

The QTL *Ae-Ps4.1,* carrying the PI180693 resistance allele, showed significant effects in the different locations and NIL genetic backgrounds studied, based on aerial and especially root data, although these have not been previously identified under controlled conditions. Leprévost et al (2023) detected this QTL only from aerial disease data in the Baccara x PI180693 RIL population. In contrast, we validated the effect of QTL *Ae-Ps5.1* in a single environment (RB) and the single DSP genetic background. This effect was previously revealed in controlled conditions with several *A. euteiches* strains from pathotypes I and III in the DSP x 90-2131 RIL population. These results suggest that the QTL *Ae-Ps5.1* may be expressed at an early stage of plant development (two week-old plants under controlled conditions) and the QTL *Ae-Ps4.1* may be expressed at a later stage (six to eight week-old plants in the field). Wang et al (2010) characterized three QTL for resistance to brown rust in barley, one acting at the seedling stage, another only from the tillering stage and the third at all stages of plant development.

The QTL *Ae-Ps2.2,* carrying the PI180693 allele, showed a significant effect on reducing aerial disease symptoms, in addition to its previously validated effect on root disease under controlled conditions. This QTL also showed a significant effect on late flowering, as identified by Hamon et al (2011). Late flowering plants stay green longer in the field, generating a bias in ADI ratings. Breaking this negative link will be essential to validate the real effect of this QTL on aerial disease symptoms.

The effect of the two QTL *Ae-Ps3.1* and *Ae-Ps 1.2* was more difficult to confirm in the NILs. At *Ae-Ps3.1*, the introgressions were lost in different NIL sets, except in NIL set 12. In this latter set, introgression of the PI180693 allele showed a significant effect on the reduction of above-ground symptoms. *Ae-Ps1.2* was previously detected with a low contribution to the phenotypic variation in the RIL population Baccara x PI180693, with an average of 5.8% and 7.3% for aerial and root disease scoring variables, respectively (Leprévost et al. 2023).

Despite the genetic x environment interactions observed, a QTL × recipient line interaction was observed for QTL *Ae-Ps7.6* from RIL BAP8.70. This resulted in an increased effect of the QTL on partial resistance in the winter pea cultivar Isard compared to the spring pea cultivars Eden, Baccara or DSP, for ADI ratings. This effect of *Ae-Ps7.6* was observed in the pea NILs under controlled conditions in Lavaud et al. (2015), from the resistance allele brought by RIL 847.50.

### QTL combinations limiting field disease severity in pea NILs

A total of four combinations of resistance alleles at two or three QTL were associated with increased partial resistance in several environments and/or NIL recipient backgrounds. These combinations included *Ae-Ps7.6* + *Ae-Ps5.1* (90-2131 alleles), *Ae-Ps7.6* + *Ae-Ps4.1* (PI180693 alleles), *Ae-Ps7.6* + *Ae-Ps2.2* ± *Ae-Ps3.1* (552 alleles) and *Ae-Ps1.2* + *Ae-Ps2.2* + *Ae-Ps3.1* (PI180693 alleles). The first three combinations were identified previously under controlled conditions (Lavaud et al. 2015).

The combination of 90-2131 alleles at the QTL *Ae-Ps5.1* and *Ae-Ps7.6* showed a significant effect on reducing symptoms on root and aerial plant parts regardless of the genetic background, with the exception of Enduro. The combination of these two QTL showed a higher effect on root areas in the spring or reference background. The combination of PI180693 alleles at QTL *Ae-Ps4.1* and *Ae-Ps7.6* showed a significant effect on root disease reduction in all three genetic backgrounds, in at least one location. It also showed a significant effect on the reduction of aerial disease symptoms in the spring variety Eden. The combination of 552 alleles at QTL *Ae-Ps7.6*, *Ae-Ps2.2* and *Ae-Ps3.1* showed a significant effect specific to Dijon and only on root symptoms in the three NIL recipient backgrounds, despite the absence of introgression at QTL *Ae-Ps3.1* in Isard and Eden. The previously unidentified combination of PI180693 alleles at the two QTL *Ae-Ps1.2* and *Ae-Ps2.2* showed a significant effect in several environments and in all three NIL recipient backgrounds. In Eden, the combination of PI180693 resistance alleles at the three QTL *Ae-Ps1.2*, *Ae-Ps2.2* and *Ae-Ps3.1*, showed a highly significant effect for both variables (DS and ADI), which was greater than that of the other combinations mentioned above. However, these allele combination at the three QTL also showed significant effect on late flowering. Hamon et al. (2013) also identified a PI180693 allelic contribution to resistance and late flowering at each of the two QTL *Ae-Ps2.2* and *Ae-Ps3.1*. Genetic linkage or pleiotropy between unwanted morphology or developmental alleles and resistance alleles should be investigated at these QTL and any possible negative linkage broken before exploiting these QTL in breeding. At *Ae-Ps2.2,* Desgroux et al. (2016) identified genetic linkage between loci associated with resistance to *A. euteiches* and flower color controlled by the *A* gene (anthocyanin production), suggesting the possibility to select peas for resistance and white flowers.

Overall, these results suggest that pyramiding QTL with strong and weak effects, as well as QTL with weak effects possibly expressed at different stages of plant development, can be promising strategies in breeding to limit Aphanomyces root rot severity. These results also suggest that the choice of QTL combinations can be guided by the geographical area of cultivation of the variety to be developed. The combination of major-effect QTL (*Ae-Ps7.6*) with minor-effect QTL possibly effective at different stages of plant development (*Ae-Ps4.1* and *Ae-Ps5.1*) and in different environnement or genetic background (*Ae-Ps1.2*, *Ae-Ps2.2*, *Ae-Ps3.1* and/or *Ae-Ps4.5*) could be relevant for selecting partially resistant varieties in regions as varied as the sites used in this study. Pyramiding these QTL could further increase the level of resistance, as demonstrated by Desgroux et al. (2016) in the AeD99OSW45-8-7 pea line, which cumulates 12 favorable haplotypes and a level of resistance much higher than that of the NILs studied carrying two or three QTL. This could also broaden the spectrum of effects of resistance on pathogen populations. In the case of bacterial downy mildew of rice, Das et al. (2022) showed the advantage of pyramiding in NILs, a major gene with three other resistance genes to improve broad-spectrum resistance while preserving yield. Pyramiding of QTL for resistance to pathogens is a promising approach to increase levels of resistance and limit QTL erosion in breeding lines (Pilet-Nayel et al. 2017).

This study identified resistance QTL combinations with effects in the field to limit root and aerial disease symptoms caused by *A. euteiches*, using NILs characterized for their introgressions and whole genomes. It suggests that pyramiding of major-effect and many minor-effect QTL is a promising strategy for breeding of resistant varieties deployable in different pea-growing regions of France. Direct use of the NILs studied and associated SNP markers in breeding schemes will help improve the resistance of pea varieties.

## Acknowledgments

We thank the INRAE experimental units of Le Rheu and Dijon-Epoisses, France, UNILET (Union Nationale Interprofessionnelle des Légumes Transformés), Quimperlé, France, for contributing to field experiments. We are grateful to Pierre Mangin and Gilles Furet for their contribution to the field experiments at Dijon-Epoisses and Chartainvilliers, respectively. We acknowledge Angélique Lesné, Isabelle Glory et Gwenola Le Roy for having contributed to DNA extraction and field trials in Riec-sur-Belon. We thank the genotyping GENTYANE platform of Clermont-Ferrand, France, for technical assistance.

## Statements & Declarations

### Funding

This work was supported by a pre-doctoral fellowship (Clément Lavaud) from INRAE, Département de Biologie et Amélioration des Plantes (France) and Brittany region (France). This work was supported by the PeaMUST project (ANR-11-BTBR-0002), which received funding from the French Government, managed by the Research National Agency (ANR) under the Investments for the Future.

### Competing Interests

The authors declare that they have no conflict of interest.

### Authors’ contributions

CL generated the phenotypic and genotypic data of the NIL, carried out statistical and genetic analyses and drafted the manuscript. JPR and PV coordinated seed preparation for all the field experiments and managed the field trials at Riec-sur-Belon et Le Rheu. GA managed the NIL genotyping using the GenoPea Infinium® BeadChip. PM, GR, JFH and PD coordinated the field trials at Dijon-Epoisses, Froissy, Mons-en-Pévèle and Louville-la-chenard, respectively. MLPN supervised the study and the draft of the manuscript. All authors approved the final draft of the manuscript.

### Availability of data and material

The datasets generated during and/or analysed during the current study are available from the corresponding author on reasonable request (Raw genotyping and phenotyping information, correlation between variables).

### Ethics approval

Authors declare that the described experiments comply with the French laws.

## ADDITIONAL FILES Legends

**Additional file 1: Detailed genomic fingerprints of pea 157 NILs at zero, one, two or three of seven main QTL for resistance to *A. euteiches* using 10 768 SNPs markers from (Tayeh et al. 2015).**

The white, dark grey, light grey, red and yellow colors indicate, respectively, for each NIL: non-polymorphic markers; polymorphic markers between the parents of the donor RILs and revealing the presence of the reference recipient genome (Baccara, Puget or DSP) in donor RILs; polymorphic markers between the parents of NILs showing a return to the recipient genome; and polymorphic markers between the parents of NILs revealing a donor fragment at homozygous or heterozygous state. Green boxes show confidence intervals of resistance QTL targeted for introgressions, according to (Leprévost et al. 2023); \**-Problem with DNA extraction and genotyping for this NIL*.

**Additional file 2: Mapping of *A. euteiches* disease scores obtained on the adjacent susceptible control (Solara) within the trials of the 157 pea NILs in contaminated nurseries, at different locations and years of experimentation.**

**A**-Disease Severity on roots; **B**-Aerial Decline Index on aerial part

**Additional file 3: Phenotypic data of disease severity on roots, aerial decline index, plant height and earliness at flowering, obtained on the 157 pea NILs and controls in each and all location*year trials studied.** LsMeans: Adjusted means; SE: Standard errors.

**Additional file 4: Mean comparison of phenotypic data for disease severity on roots, aerial decline index, plant height and earliness at flowering obtained on all trials, by NIL set.**

LsMeans: Adjusted means; The different letters represent significant differences between NILs (Tukey-HSD test, α=5%).

**Additional file 5: Mean contrast values between NILs with and without QTL, within each NIL set for disease severity on roots (5a) and aerial decline index (5b) in each and all location*year trials**

A color scale from red to green represents a resistance level of a QTL-carrying NIL from lower to higher, respectively, than that of the QTL-free NIL from the same set. In bold: Significant differences of contrast values between the QTL-carrying NILs and the average values of the two sister NILs without QTL; NA: missing data.

**Additional file 6: Description of interactions between QTL, individually or in combination, and the genetic background, for disease severity on roots (DS) and aerial decline index (ADI), by donor NIL parental line.** The letters represent significant differences between NILs carrying the same QTL, individually or in combination, in the different genetic backgrounds (Tukey-HSD test, α=5%).

## Notes

### Competing Interest Statement

The authors have declared no competing interest.

