## Supplementary figures and images for "QTL combinations associated with field partial resistance to aphanomyces root rot in pea Near-Isogenic Lines"

### Additional file 2

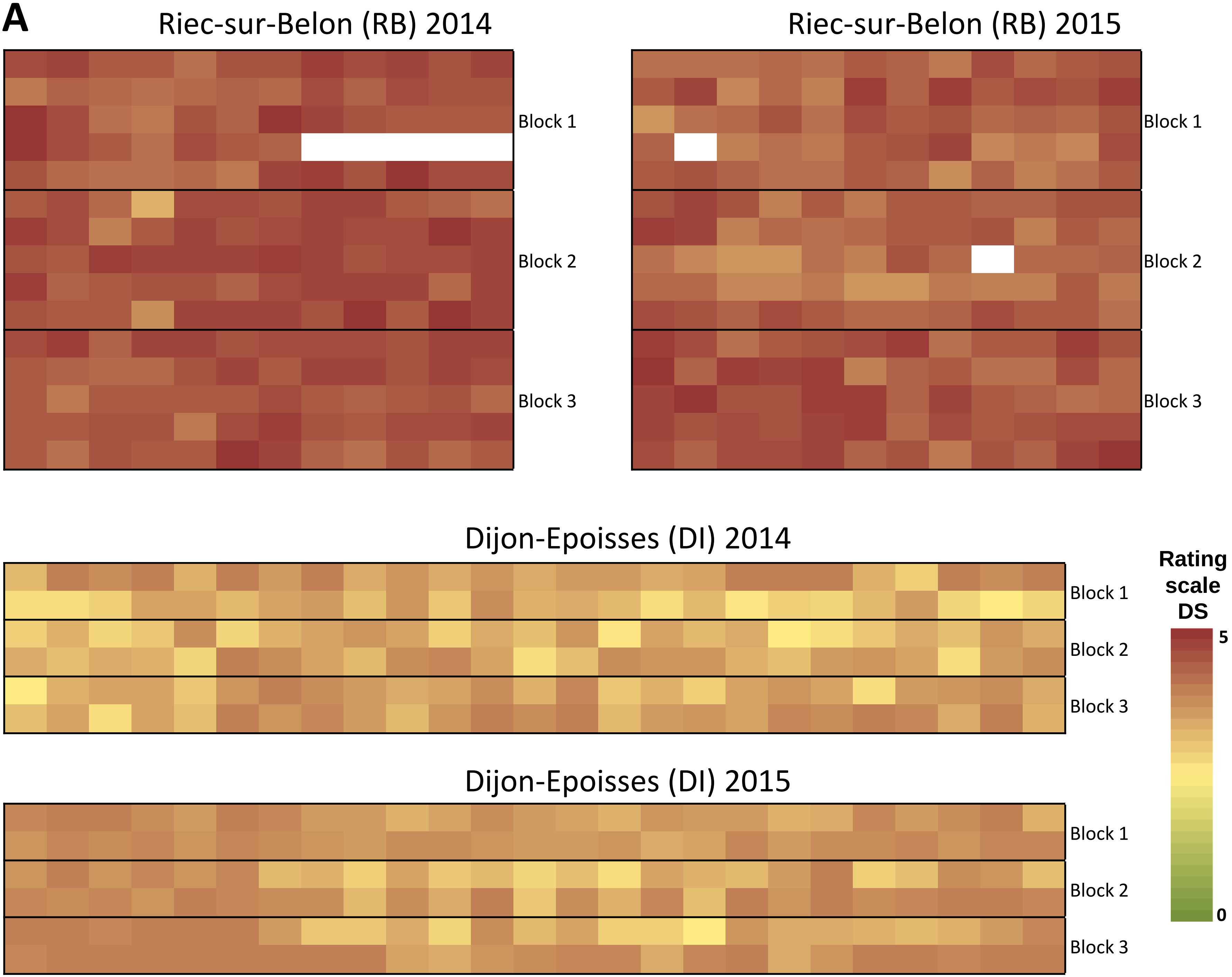

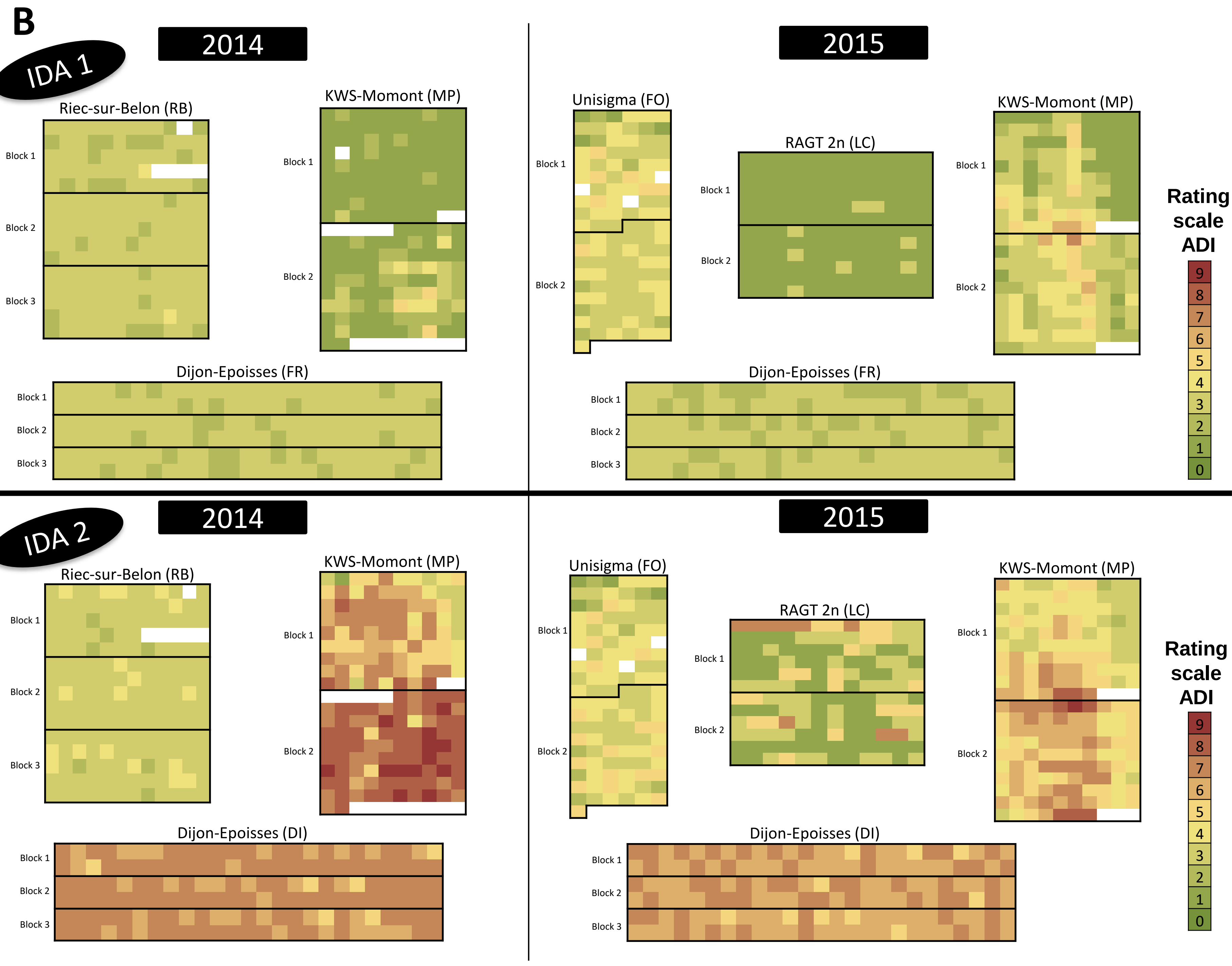

### Additional file 6

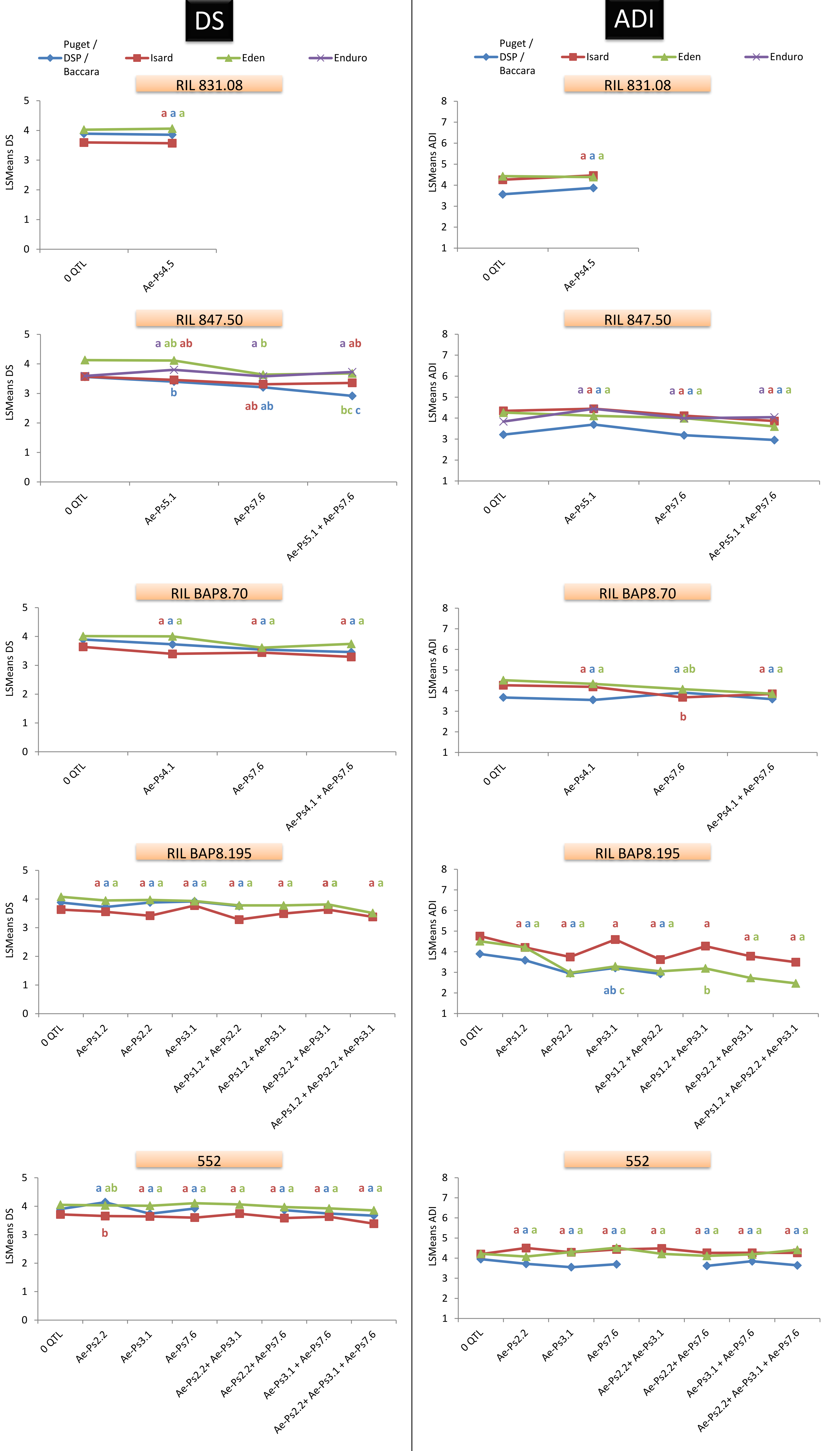
